# Stereoselective binding of prasugrel active metabolite to the P2Y12 receptor: insights from a molecular modeling approach

**DOI:** 10.64898/2026.03.26.713933

**Authors:** Florentin Allemand, Laura Le Bras, Siamak Davani, Christophe Ramseyer, Jennifer Lagoutte-Renosi

**Affiliations:** Université Marie et Louis Pasteur, SINERGIES, F-25000 Besançon, France; Université Marie et Louis Pasteur, CNRS, Chrono-environnement, F-25000 Besançon, France; Laboratoire de Biologie Structurale de la Cellule (BIOC), CNRS, École polytechnique, Institut Polytechnique de Paris, 91120 Palaiseau, France; Université Marie et Louis Pasteur, CHU Besançon, SINERGIES, Service de Pharmacologie Clinique et Toxicologie, F-25000 Besançon, France

## Abstract

Prasugrel is a prodrug, widely used in antiplatelet strategy for secondary prevention after acute coronary syndrome. The metabolism of prasugrel leads to the formation of the Prasugrel Active Metabolite (PAM), an irreversible P2Y12 receptor antagonist. Its mode of binding has not yet been fully established, although it is known that it binds covalently to P2Y12 by forming a disulfide bridge with cysteines and its sulfur moiety. PAM is a molecule with two chiral centers, resulting in four stereoisomers which appear to be stereoselective upon binding. A combination of different molecular modeling methods, such as molecular dynamics, ensemble docking, and Density Functional Theory (DFT), were used to rationalize these differences in antagonism observed *in vitro* and to elucidate the mode of binding of PAM to P2Y12. PAM is found to bind to the closed P2Y12 conformation in a preferential way. Although the four stereoisomers have comparable affinity, the location of the RS stereoisomer makes the formation of a disulfide bond with cysteines more favorable, particularly with cysteine 175. Compared to the RR stereoisomer, the RS stereoisomer interacts less deeply with the P2Y12 receptor, interacting in particular with the second and third extracellular loops, explaining the competition observed with cangrelor and an intermediate metabolite of prasugrel. Furthermore, DFT calculations have shown that the formation of a disulfide bridge is energetically more favorable with the RS stereoisomer than with the RR stereoisomer. The physical interactions and chemical reaction between the RS stereoisomer and the P2Y12 receptor are key factors in explaining the stereoselective binding of PAM to P2Y12.

## 1 Introduction

Antiplatelet therapy is indicated in all acute coronary syndrome (ACS) patients to prevent secondary cardiovascular events. The therapeutic strategy consists nowadays of a combination of two antiplatelet agents, combining a P2Y12 receptor antagonist and aspirin as a cyclooxygenase (COX) inhibitor according to the European Society of Cardiology ^1^. Regarding P2Y12 receptor for Adenosine DiPhosphate (ADP), two classes of antagonists are available on the market, namely irreversible thienopyridines and reversible ADP analogues. Thienopyridines, such as ticlopidine, clopidogrel, and prasugrel, are prodrugs metabolized by the liver. Their active metabolite covalently binds to a cysteine residue on the P2Y12 receptor, thereby preventing ADP binding ^2^. Regarding prasugrel, its recognition by P2Y12 remains unfortunately poorly understood at the molecular level. In contrast to this strong chemical bond, reversible P2Y12 receptor antagonists such as ticagrelor and cangrelor have been designed to mimic the physical binding of ADP with weaker interaction energies.

In this work, we focused more on the binding of prasugrel using molecular simulations. The first reason is because prasugrel and ticagrelor are recommended in preference to clopidogrel. Moreover, prasugrel should be considered in preference to ticagrelor for ACS patients who undergo percutaneous coronary intervention ^1^. The second reason is that the precise mode of binding of prasugrel is unknown, unlike Ticagrelor, where we only know that cysteine 97 or 175 is involved ^3^.

Concerning the ligand, the Prasugrel Active Metabolite R-138727 (PAM) is known to form a disulfide bridge with cysteine residues of P2Y12. It has been demonstrated that cysteines 97 or 175 are involved in this binding process ^4,5^. Algaier *et al.* showed that if one of these cysteines is mutated, PAM no longer inhibits the P2Y12 response to an agonist ^4^. Interestingly, a disulfide bridge exists between these two cysteines, but it is labile ^6^ allowing the ligand to break this bridge and create its own bond with one of the cysteine residue. Furthermore, from a chemical point of view, PAM has two chiral centers, compared with one for prasugrel. The metabolization of prasugrel to form PAM induces the creation of a new chiral center, giving rise to four possible stereoisomers, namely RS, SR, RR, and SS., Hasegawa *et al.*, measured the platelet aggregation induced by ADP, as a function of the concentration of each of these stereoisomers ^7^. They revealed that stereoselectity played an essential role. They showed that RS stereoisomer (IC_50_: 0.19 µM) has the higher potency in inhibiting ADP-induced platelet aggregation, followed by RR stereoisomer (IC_50_: 3.1 µM), and finally SS (IC_50_: 28 µM) and RR (IC_50_: 36 µM) which have similar inhibition. The binding of PAM to P2Y12 therefore appears to be stereoselective, with RS stereoisomer being significantly superior. However, no clear explanation was given to explain this selectivity. Additionally, the formation of the active metabolite by cytochromes in the liver is a stereoselective process. Wickremsinhe *et al.* have shown that the RS and RR stereoisomers represent 84%, each corresponding to about 42%, of the metabolites present in the plasma after administration of prasugrel, compared to only 16% for the SS and SR stereoisomers ^8^. Therefore, it appears that the two most active geometries are produced in the greatest quantity.

Concerning the P2Y12 receptor, there are also some bottlenecks that render the determination of the binding process difficult. The main one concerns the P2Y12 structure and dynamics. P2Y12 is a G Protein-Coupled Receptor (GPCR) embedded in the platelet plasma membrane and that exhibits several conformations upon binding ligands. Two possible conformations of the P2Y12 receptor, namely an open and a closed one have been elucidated ^6,9,10^. The difference between these two conformations lies mainly in the conformation of the sixth transmembrane helix. In the closed conformation, observed in the presence of an agonist, an angle appears in the upper third of the helix, closing the helix towards the inside of the receptor whereas in the open conformation, observed in the presence of an antagonist, the helix is straight. These two conformations differentially bind P2Y12 agonists and antagonists, but the specific interactions of PAM with these two conformations have not yet been studied. The second reason that makes the investigation of P2Y12 difficult is its immediate environment in the membrane. Indeed, this receptor, like many other GPCRs, is very sensitive to its lipid environment ^10,11^. Lipidomics study of platelet plasma membrane revealed that it contains high levels of arachidonic acid, a 20:4 polyunsaturated fatty acids ^12^. In addition, the platelet plasma membrane contains domains rich in arachidonic acid where P2Y12 receptors are preferentially localized ^13^. It is therefore important to take into account this local environment containing arachidonic acid.

The aims of this study are firstly to elucidate the binding mode of PAM to P2Y12 and secondly to rationalize the difference in *in vitro* antagonism of these stereoisomers. It seems important to understand the stereoselectivity of P2Y12 with respect to PAM in order to optimize the targeting of P2Y12 receptor by future antiplatelet agents.

To this end, Molecular Dynamics (MD) calculations were carried out to determine the dynamics of the P2Y12 receptor in open and closed conformation in its lipid environment, and to study the interactions of stereoisomers bound to P2Y12. An ensemble docking approach was used to quantify and determine the binding modes of the four stereoisomers on the P2Y12 receptor.

Finally, quantum chemistry calculations using Density functional Theory (DFT) were carried out to quantify the formation of a disulfide bridge between PAM and P2Y12.

## 2 Materials and methods

### 2.1 Molecular Dynamics

MD was performed with GROMACS 2018 ^14^ to simulate the dynamics of the P2Y12 receptor in a realistic lipid environment. The CHARMM36 force fields were used for the protein and lipids ^15,16^. We considered the two conformations which have been crystallized ^6,9^. The open (4NTJ) and closed (4PXZ) forms of P2Y12 were inserted within a lipid bilayer of around 400 lipids built with CHARMM-GUI ^17,18^. The composition of this bilayer was established from experimental data on the lipid composition of the plasma membrane of platelets (Table 1) ^12^. These different lipids were distributed between the outer and inner leaflets, knowing that the SphingoMyelins (SMs) are located in the outer leaflet while the PhosphatidylSerines (PSs) and the majority of the PhosphatidylEthanlolamines (PEs) are in the inner leaflet ^19^. Regarding the PEs, the platelet plasma membrane mainly contains their plasmalogen form, which are here OAPE and YAPE. A cholesterol amount of 30% was added in accordance with lipidomic data ^12^. Sodium and chloride ions were added to achieve isotonic blood plasma conditions. All systems were hydrated to approximately 75 water molecules per lipid head. The systems were first equilibrated for 250 ps under NVT conditions with the Berendsen thermostat. Then, they were equilibrated under NPT conditions, with the Berendsen thermostat and barostat ^20^, for 200 ns, divided into two phases of 100 ns each. First, the system was equilibrated by constraining the protein with a force constant of k = 1000 kJ mol^-1^ nm^2^, then only its backbone was constrained during the second phase. A 200 ns production phase was then simulated.

**Table 1.** Lipid composition of the bilayer for molecular dynamics simulations of P2Y12 in its lipid environment. CHOL = Cholesterol. Phospholipids are composed of two chains of fatty acids in sn1 or sn2 position: M =Myristic, L = Linoleic, P = Palmitic, O = Oleic, A = Arachidonic, S = Stearic, Y = Palmitoleic. They also contain a specific polar head: PC = PhosphatidylCholine, PE = PhosphatidylEthanolamine, PS = PhosphatidylSerine, SM = SphyngoMyelin. There is an asymmetry between the outer (first line) and inner leaflet (second line)

| CHOL | MLPC | PLPC | POPC | PAPC | SAPE | OAPE | YAPE | SAPS | PSM | BSM | Total |
| --- | --- | --- | --- | --- | --- | --- | --- | --- | --- | --- | --- |
| - | 14:0 | 16:0 | 16:0 | 16:0 | 18:0 | 18:2 | 16:2 | 18:0 | 18:1 | 18:1 |  |
| - | 18:2 | 18:2 | 18:1 | 20:4 | 20:4 | 20:4 | 20:4 | 20:4 | 16:0 | 22:0 |  |
| 75 | 25 | 12 | 37 | 8 | 4 | 4 | 4 | 0 | 16 | 16 | 211 |
| 48 | 16 | 8 | 24 | 6 | 24 | 20 | 16 | 28 | 0 | 0 | 198 |

Another set of MD simulations was done with the closed conformation of the P2Y12 receptor to which the RS and RR stereoisomers are covalently bound to CYS97 and CYS175, the two cysteines with which PAM can bind ^4^, in order to study the binding site of these two stereoisomers. Results of covalent docking calculations performed with AutoDock ^21^ between sulfur atom from cysteine 175 of the P2Y12 receptor and sulfur atoms from these two stereoisomers were used to build these models.

### 2.2 Ensemble docking

Docking calculations of PAM stereoisomers were conducted on both P2Y12 conformations. Ensemble docking is a method that allows protein dynamics to be considered, as docking is not done on a single specific conformation of the protein in this case but on a wide range of conformations that the protein adopts over a few tens of nanoseconds. The second advantage of this procedure is that it allows statistics to be determined on a large set of PAM/P2Y12 binding poses and scores. In this work, the last 50 nanoseconds of MD simulations were used. A single structure was extracted every 60 ps, meaning that 833 conformations of P2Y12 were selected for docking. The docking of the four stereoisomers of PAM on these structures was carried out using quickvina-2 ^22^. The carboxylic acid function of PAM was deprotonated for these calculations in agreement with its pKa ^23^. The docking box was defined by a cube with sides of 2 nm, centered on the active site of P2Y12 ^6,9^. The best poses from each docking calculation were analyzed in order to study the interactions between P2Y12 and the stereoisomers of PAM using MDAnalysis ^24,25^.

### 2.3 Quantum chemistry

The chemical interaction between PAM and P2Y12 can be investigated with quantum chemistry calculations which are well-suited to describe the bonds formation. Density Functionnal Theory (DFT) calculations were thus performed in this work to energetically study the disulfide bond formation between cysteine 175 of the P2Y12 receptor and the RS and RR stereoisomers. This cysteine was chosen, rather than cysteine 97, based on ensemble docking results (see section 3). These models were prepared using the results of covalent docking calculations performed with AutoDock ^21^ between the sulfur atom of cysteine 175 and the sulfur atoms of these stereoisomers. Models were then hydrated using AutoDockTools ^26^, and a minimization step was performed to relax the position of these hydrogen atoms. From these structures, a binding site was isolated and then passivated by adding hydrogen where the protein structure has been cut using the quantum chemical clustering approach (Figure 1) ^27^. In order to maintain the protein’s structure during quantum calculations, the position of the alpha carbons of all residues was frozen. QM/QM’ calculations were performed with ONIOM approach from Gaussian09 software ^28^. B3LYP/6-31g(d) exchange-correlation functionnal/basis set combination was used for the low level model and MPW1N/6-31g(d) for the high level model, which contains the two sidechains of the two cysteines and the sulfur and bound carbon from PAM. MPW1N is particularly well suited for describing Thiol−Disulfide Exchange ^29,30^. An optimization calculation was first performed. Next, a frequency calculation was performed to assess the energy of the optimized structure and check that the structure was in a true minimum. The transition state was determined by placing the sulfur atom of cysteine 175 at equal distance from those of PAM and cysteine 97. The optimized geometry of the transition state was validated by a frequency calculation highlighting a unique imaginary frequency corresponding to the exchange of the disulfide bond.

**Figure 1.**
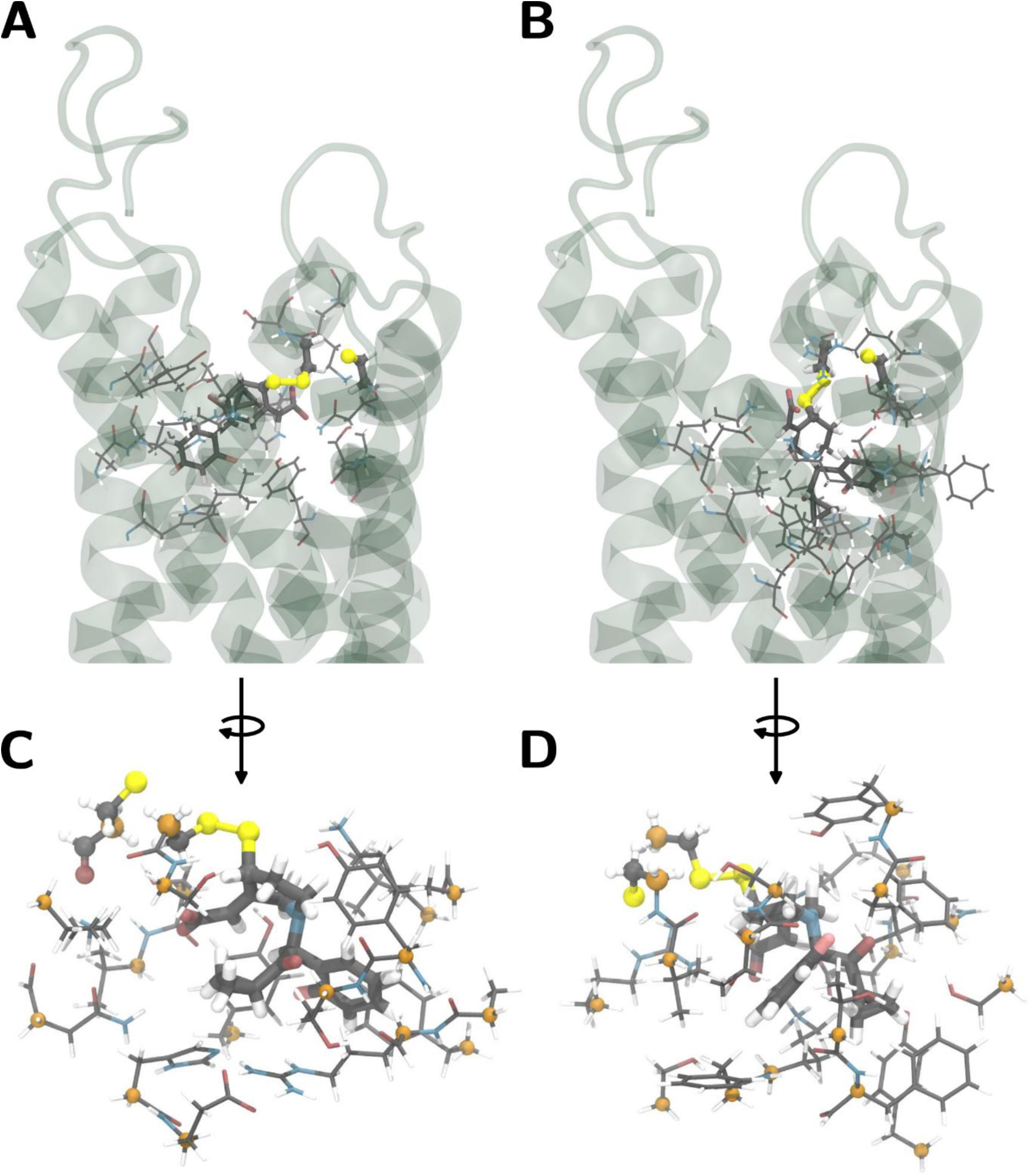
**Models of the binding sites of RS (A) and RR (B) stereoisomers bind to cysteine 175 simulated by QM/QM’ approach** represented with the structure of the P2Y12 receptor and without (D-C). Hydrogen atoms attached to carbons are not shown. P2Y12’s residues are shown as sticks, PAM stereoisomers as licorice and the high model, including cysteines 97 & 175 and thiol function from PAM, as balls and large licorice. The alpha carbons frozen during DFT calculations are shown in orange. The other carbon atoms are shown in gray, hydrogen in white, oxygen in red, nitrogen in blue and sulfur in yellow.

## 3 Results

As mentioned above, we choose to decompose the PAM/P2Y12 interaction in a two steps process governed by physical (Figures 2-4) and chemical interactions (Figure 5), respectively. Figure 2 represents the distribution of the best affinities resulting from ensemble docking calculations on P2Y12 open and closed conformations.

**Figure 2.**
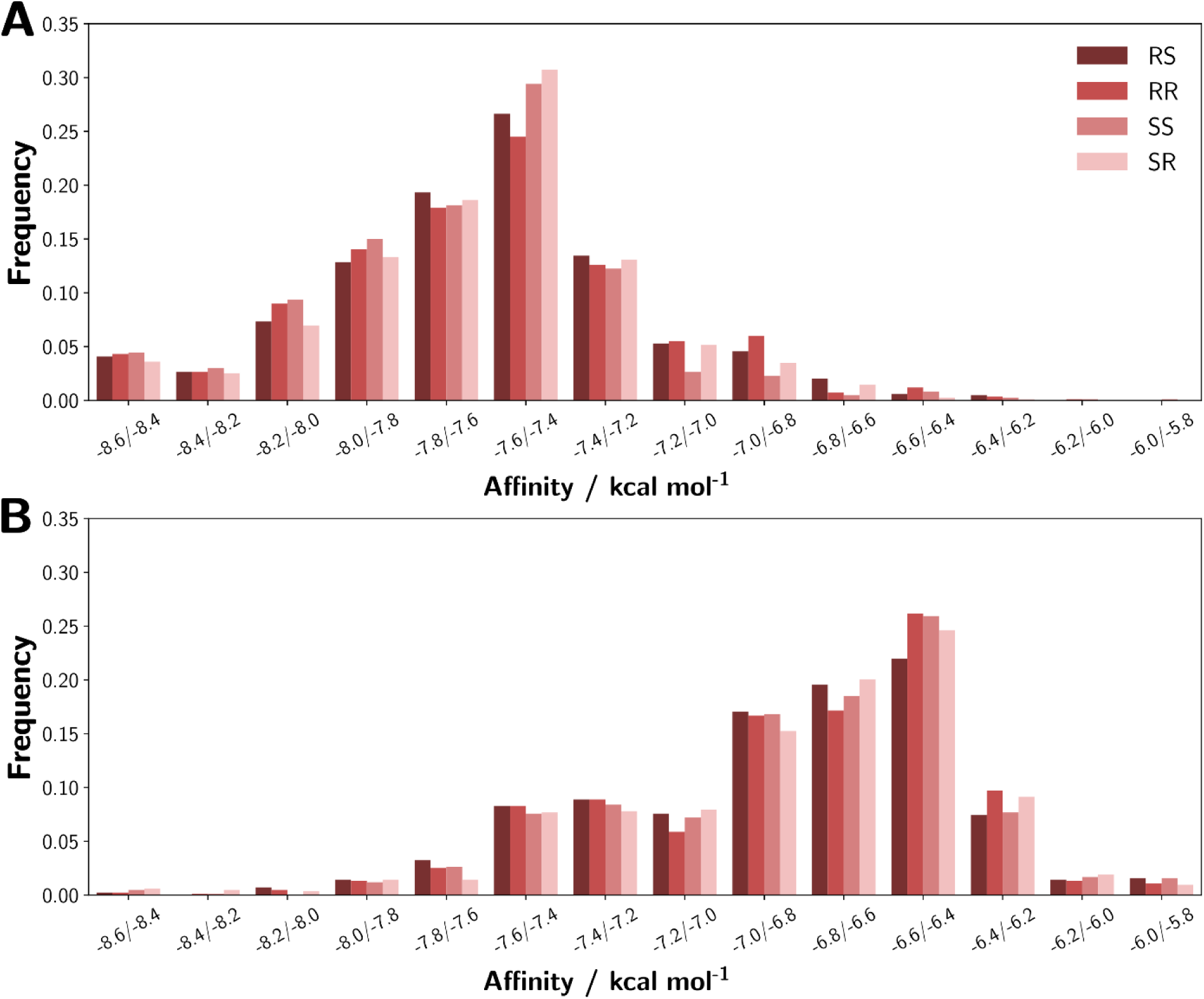
**Distribution of affinities from ensemble docking of the four Prasugrel Active Metabolite (PAM) stereoisomers for P2Y12** in (A) Closed and (B) Open conformations.

The closed conformation appears to be more favorable compared to the open conformation, regardless of the stereoisomer. For the closed conformation (Figure 2A), there is no difference in frequency of occurrence between the two most favorable affinity classes, from -8.6 to -8.2 kcal mol^-1^. For the next two classes, from -8.2 to -7.8 kcal mol^-1^, the RR and SS stereoisomers are slightly overrepresented. For the fifth most favorable class, from -7.8 to -7.6 kcal mol^-1^, the RS stereoisomer is more represented. Regarding the open conformation (Figure 2B), the -7.8 to -7.6 kcal mol^-1^ class highlights an overrepresentation of the RS stereoisomer and an underrepresentation of the SR stereoisomer. For the next two classes, from -7.6 to -7.2 kcal mol^-1^, the RS and RR stereoisomers are more represented than the other two. Overall, the open conformation is more favorable to the RS, followed by the RR stereoisomer, compared to the SR and SS stereoisomers. At this stage, it should be noted that PAM forms a covalent bond with cysteine from P2Y12 via a disulfide bridge with a typical length of 2.04 Å ^31^. The sulfur atoms of the ligand and the receptor must be close enough to allow the thiol-disulfide exchange. Since disulfide bridges are suspected to play a key role in that case, the distances between the sulfur atom of PAM and that of cysteines 97 and 175 were analyzed (Figure 3).

**Figure 3.**
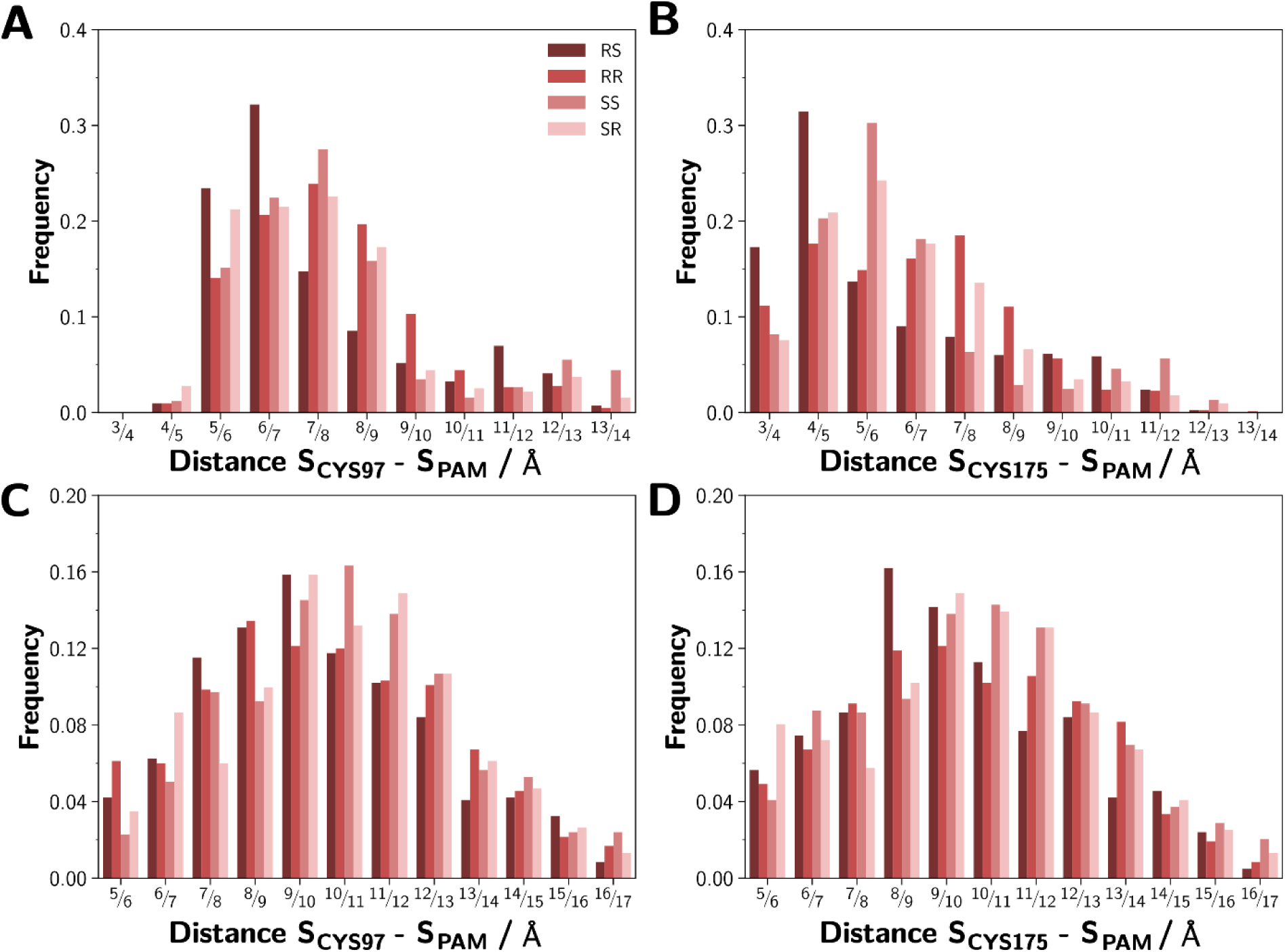
**Distribution of S-S distance between cysteines 97 (A & C) and 175 (B & D) and the thiol of the four Prasugrel Active Metabolite (PAM) stereoisomers** for poses from ensemble docking on P2Y12 in closed (A & B) and open (C & D) conformation.

Figure 3 presents the distribution of distances between the sulfur atom of these residues and the sulfur atom from the thiol group of the different stereoisomers. They were extracted from ensemble docking poses obtained on the last 50 ns of MD simulations. For the closed conformation, the most common shortest distances are between the RS stereoisomer and cysteine 175, with nearly 50% of the poses resulting from ensemble docking showing a distance less than 5 Å, following by the RR stereoisomer. The distances are always longer with cysteine 97 compared to cysteine 175, but the RS stereoisomer is the only one with more than 50% of distances less than 7 Å with cysteine 97. For the open conformation, the distances are on average greater than for the closed conformation. Moreover, the distribution is more homogenous without marked difference between the two cysteines in this open conformation, unlike what is observed with the closed conformation. However, the RS stereoisomer shows distances slightly shifted towards the 7 to 9 Å range compared to the other stereoisomers. Taken together, these results suggest that the RS stereoisomer is the one with the greatest propensity to establish a disulfide bridge with P2Y12, in closed conformation, and more particularly with cysteine 175.

In a second step, we tried to identify the binding site of the two stereoisomers, RS and RR, with the greatest antagonism for P2Y12. This approach was based on the results of experiments carried out by Algaier *et al.* who showed that PAM could bind to cysteine 97 and to cysteine 175 via a disulfide bond ^4^. According to the structure of P2Y12, these two residues are very close, being also linked by a disulfide bridge ^6,9^. We assume that ligands should be able to exchange this disulfide bridge to form another, either with cysteines 97 or 175. Analysis of the contacts between RS and RR stereoisomers, when they are covalently bound to cysteine 97 and cysteine 175, and P2Y12 receptor residues reveal the binding pockets of these two molecules (Figure 4). Overall, the binding pocket of the different ligands is stable during the 200 ns simulations, as persistent contacts are present for each of the stereoisomers. These contacts involve many charged residues or residues capable of forming hydrogen bonds in contact with the two stereoisomers. This is particularly true for residues in contact with both stereoisomers bound to cysteine 97 for most of the simulation time: arginine 93 and 256, histidine 187 and threonine 260. These four residues are also in contact with the RS stereoisomer when it is bound to cysteine 175, but not with the RR one. The RR stereoisomer bound to cysteine 175 has different contacts, but also with charged residues such as lysines 80 and 280 or those that can form hydrogen bonds such as tyrosines 32 and 105, threonines 76 and 100, and serines 83, 101 and 288. Other residues are more specific to one of the two stereoisomers. The RR stereoisomer bound to cysteine 97 interacts with residues from helices III, IV, V and VI, while the RS stereoisomer interacts with residues from helices VI and VII. In addition, both interact with the second extracellular loop, but with different residues. For example, the RS stereoisomer interacts with the third extracellular loop, while the RR stereoisomer does not. Overall, the RR stereoisomer interacts with deeper residues within the receptor, while the RS stereoisomer remains closer to the surface. This is also observed at the end of the simulation, where RR, whether bound to cysteine 97 or 175, is deeper than RS (Figure 4 B, C, D & E).

**Figure 4.**
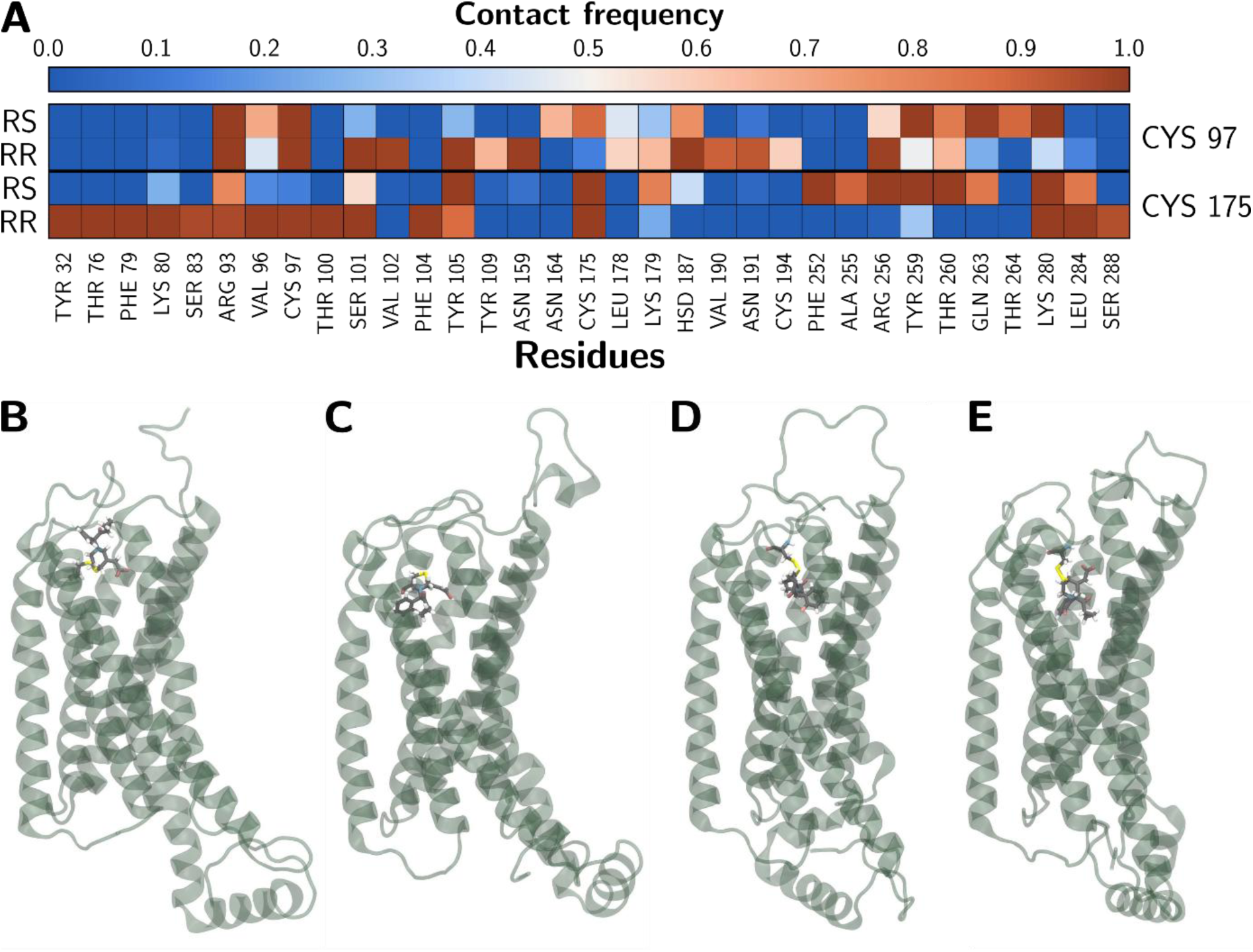
Frequency of contact between RR and RS stereoisomers with different P2Y12 residues (A) and localization of PAM stereoisomers at the end of simulation (B-E). The frequency of contact was calculated over the whole 200 ns production phase of simulations where stereoisomers are covalently bound to cysteines 97 and 175 of P2Y12. Contact is defined as a distance of less than or equal to 4 Å between a stereoisomer atom and a residue atom. Only residues with a contact frequency of at least 0.5 with at least one of the two stereoisomers and cysteines are shown. Limits of helices: Helix I: 26-51, Helix II: 57-83, Helix III: 96-122, Helix IV: 139-164, Helix V: 188-214, Helix VI: 233-260, Helix VII: 278-304. P2Y12 is oriented with its V^th^ helix in the foreground with RS (B) and RR (C) stereoisomers bind to cysteine 97 and RS (D) and RR (E) stereoisomers bind to cysteine 175.

Finally, DFT calculations have made it possible to energetically quantify the chemical reaction that allows to break the disulfide bridge between the cysteines 97-175 and create the bond between cysteine 175 and PAM (Figure 5). This reaction has an energy barrier of 15.5 kcal mol^-1^ for the RS stereoisomer and a higher energy barrier of 25.3 kcal mol^-1^ for the RR stereoisomer. For the RS stereoisomer the Gibbs free energy of reaction is Δ_r_G_RS_ = 12.1 kcal mol^-1^ and Δ_r_G_RR_ = 10.8 kcal mol^-1^ for RR stereoisomer. These reactions are endergonic which means that additional energy needs to be provided in order to promote the reaction. It should be noted that it has a lower energy than the spontaneous dissociation of a disulfide bridge (60 kcal mol^-1^) ^32^ which can occur naturally since this disulfide bridge is labile. This higher barrier for the RR stereoisomer may partly explain the difference in aggregation inhibition observed *in vitro*. The key parameters appear to be the energy required for the formation of the bond itself and the localization of PAM when it interacts with P2Y12. A position that further promotes the formation of this bond in the case of the RS stereoisomer, due to the shorter distances between the sulfur atom of the ligand and that of cysteine 175.

**Figure 5.**
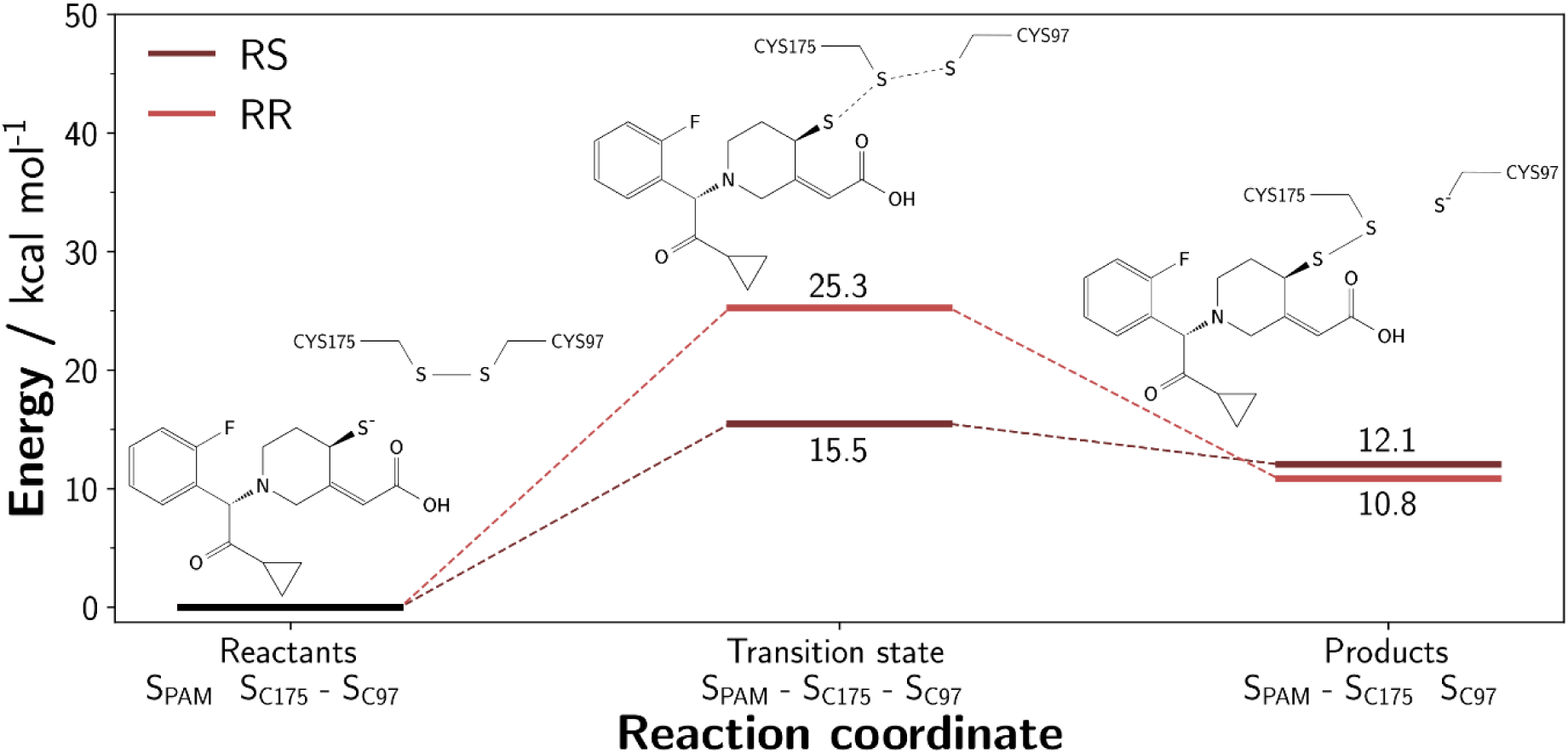
**Energy profile of the disulfide bond exchange** between cysteines 97 and 175 and cysteine 175 and the RS and RR stereoisomer of Prasugrel Active Metabolite (PAM).

## 4 Discussion

This study provides the first molecular-level characterization of the binding mode of the prasugrel active metabolite with its target, the P2Y12 receptor. We obtained four main results in that work that may be helpful for design of new covalent P2Y12 inhibitors. First, we established that PAM prefers to bind into the closed state conformation of P2Y12. Second, we showed that the RS stereoisomer, followed by the RR one, has the better propensity to establish a disulfide bridge with P2Y12 which could explain its most potent antagonism for P2Y12 compared to other stereoisomers. Third, we determined that cysteine 175 is the closest to the thiol function of PAM stereoisomers, making it possible for a disulfide bridge to form between them. Fourth, we observed that the RS stereoisomer remains closer to the surface of P2Y12 receptor while the RR stereoisomer interacts with deeper residues within P2Y12 receptor. Fifth, DFT calculations have shown that the formation of a disulfide bridge is energetically more favorable with the RS stereoisomer than with the RR stereoisomer.

Regarding the difference in antagonism between the four stereoisomers observed *in vitro*, the affinities obtained by ensemble docking do not explain it. Indeed, the distribution of non-covalent affinities for P2Y12, in open and closed conformations, scored by ensemble docking is consistent for all four stereoisomers. It appears that interactions with the closed conformation are always more favorable than those with the open conformation. This is consistent with the fact that the closed conformation is known to be the active form and the open conformation the basal form of the P2Y12 receptor ^33^. Hence, PAM preferentially binds to the active form of the P2Y12 receptor. The same applies to another P2Y12 antagonist, ticagrelor, and to the agonist, ADP ^10^.

The interaction of the RS stereoisomer with P2Y12 in the closed conformation does not appear to be more favorable than that of the other stereoisomers. Consequently, this cannot explain why the RS stereoisomer is the one most likely to bind to P2Y12 via a disulfide bridge according to *in vitro* experiments ^7^. This disulfide bridge forms after a thiol-disulfide exchange with the existing disulfide bridge between cysteines 97 and 175. The thiol of the ligand must therefore be located close to this disulfide bridge. For the closed conformation of P2Y12, which is most favorable for PAM binding, the RS stereoisomer, compared to the other stereoisomers, has its sulfur atom closest to the sulfur atoms of the two cysteines. In particular, it is closer to cysteine 175 than to cysteine 97. After the RS stereoisomer, RR has the shortest distances, followed by SS and finally SR. This corresponds to the ranking in order of potency in inhibiting ADP-induced aggregation by these four stereoisomers ^7^. Similarly, Deflorian *et al.* showed that the sulfur atom of the clopidogrel active metabolite, of which prasugrel is an analog, is close to that of cysteine 175 (4.7 Å) when it binds to the P2Y12 receptor ^34^.

If the sulfur-sulfur distances between the RS and RR stereoisomers and cysteine 175 are different, this can be explained by the fact that their binding sites are different. Indeed, we demonstrated that RS and RR stereoisomers do not interact with the same P2Y12 residues when the RS and RR stereoisomers are covalently bound to cysteine 97 or 175. The RR stereoisomer is located deeper within the receptor than the RS one, as it makes contacts mainly with transmembrane helix residues, whereas the RS stereoisomer mainly interacts with extracellular loops, notably the second extracellular loop. Minuz *et al.* have shown that a Prasugrel Intermediate Metabolite (PIM) can bind to P2Y12 in a reversible way, particularly at this second extracellular loop ^35^. They also showed that this PIM negatively interferes with PAM, thereby reducing its antagonism. Competition between PAM and PIM appears to occur at the level of the RS stereoisomer binding pocket. It is interesting to note that cysteine 175 is located within this second extracellular loop. Thanks to our study, we can assume that PIM could prevent PAM from covalently binding to cysteine 175, reducing the effectiveness of PAM on P2Y12. In addition, Judge *et al.* showed *in vitro* that cangrelor, another P2Y12 antagonist, prevented PAM from binding to P2Y12 ^36^. Cysteine 175 is found in the residues forming the cangrelor binding pocket, but also several residues in contact with PAM covalently bound to cysteine 175 are found in this pocket: Arg93, Tyr105, Phe252, Tyr259, Gln263, Lys179, Arg256, and Lys280 (Figure 4) ^9,37^. Taken together, these results support that RS stereoisomer of PAM covalently binds to P2Y12 in closed conformation via cysteine 175 rather than via cysteine 97 (Figure 6).

**Figure 6.**
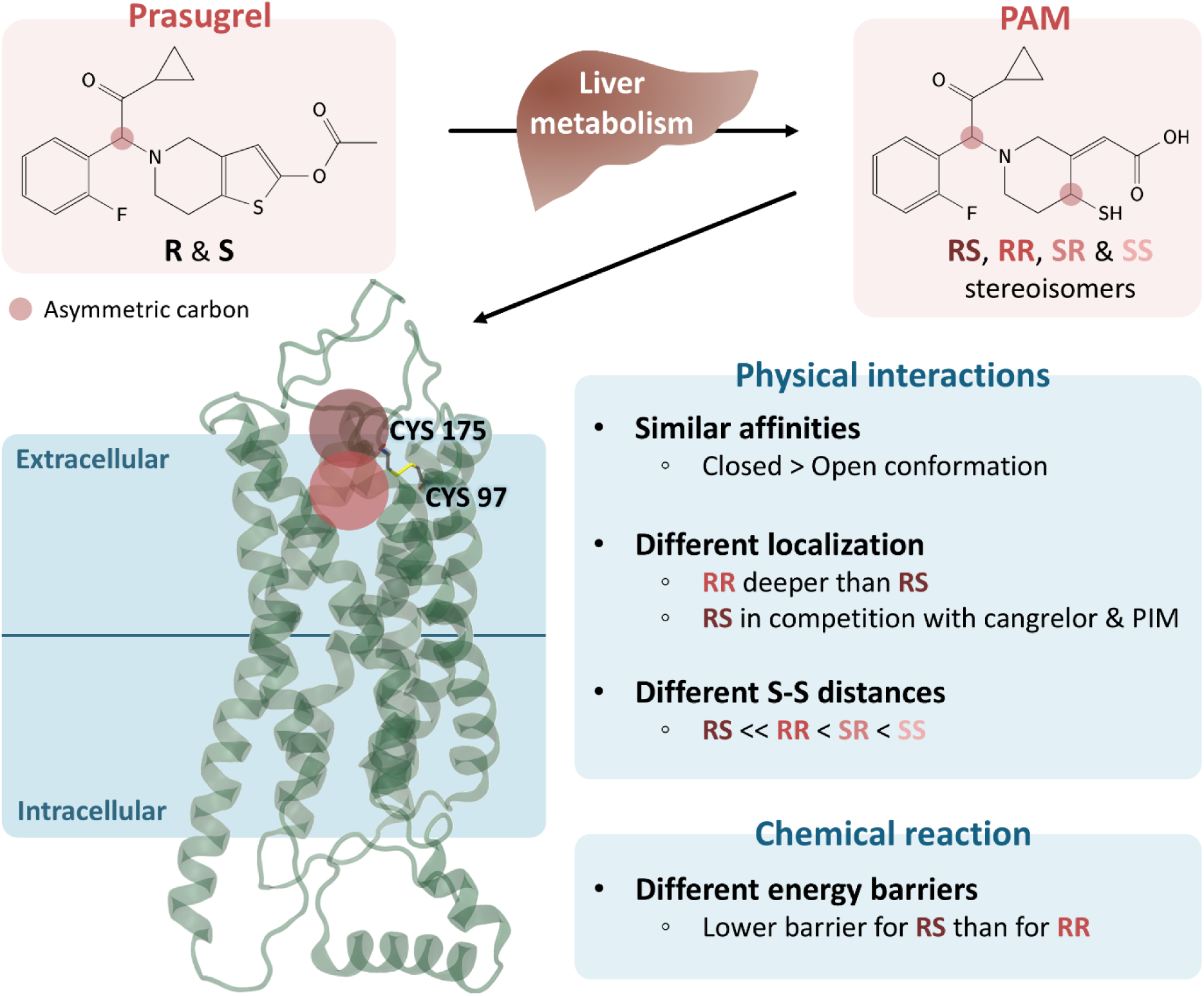
Schematic representation of the conclusions concerning the binding of different Prasugrel Active Metabolite (PAM) stereoisomers to P2Y12. The distance between the sulfur atom of the ligand and the cysteines 97 & 175 of P2Y12 is shortest with the RS stereoisomer and cysteine 175. The RS stereoisomer binds less deeply than the RR stereoisomer and competes with cangrelor and an Prasugrel Intermediate Metabolite (PIM).

Regarding the study of the formation of the bond between RS and RR stereoisomers and cysteine 175, we showed that both reactions are endergonic, but with a lower energy barrier for the RS stereoisomer. Furthermore, the existing disulfide bridge between cysteine 97 and cysteine 175 is labile ^6^, so the formation of a disulfide bond between PAM and cysteine 175 could occur from a state where cysteine 175 is free. This could be more favorable for the formation of this disulfide bond. The activation energies of RS and RR stereoisomers bond-formation at cysteine 175 differ, unlike their Gibbs free energies, which means that bond formation with the RS stereoisomer is more favorable. In addition, as explained above, the interactions of RR and RS stereoisomers with P2Y12 involve different locations, deeper for RR and more superficial for RS. As a result, the RS stereoisomer is in a more favorable position to form a disulfide bond with cysteine 175, due to the short distances between its sulfur atom and that of cysteine. This tends to show that chemical reaction and the physical interactions of PAM within the P2Y12 binding site are complementary key steps in the binding process. It is probably these steps that enable the RS stereoisomer to have greater antagonistic effect on P2Y12 receptor. This paves the way for the development of new antiplatelet agents targeting this disulfide bond.

## 5 Conclusions

The binding of PAM to the P2Y12 receptor is stereoselective given stereoisomers inhibit platelet aggregation differently. Ensemble docking simulations revealed a similar affinity of the four stereoisomers of PAM for the P21Y2 receptor in its closed conformation. However, these similar affinities do not imply the same interactions with the receptor. We showed that the RS stereoisomer localizes within the P2Y12 receptor by presenting its sulfur atom close to that of cysteine 175, making disulfide bond formation more favorable than the other stereoisomers. Additionally, RS stereoisomer is located less deep than the RS stereoisomer within P2Y12 receptor, and in the same binding pocket as other antagonists which competitively bind to the P2Y12 receptor. The study of bond formation also shows that it is energetically more favorable with the RS stereoisomer. Overall, the conformation of the RS stereoisomer and the energy required to form a disulfide bridge with cysteine 175 are the key factors explaining its greater ability to form a covalent bond with the P2Y12 receptor.

